# AI-Driven Identification and Validation of RXFP1 as a CML Biomarker Using Gene Expression and Integrated GUI Tools

**DOI:** 10.1101/2025.06.06.658261

**Authors:** Mohd Sufiyan Khan

## Abstract

Chronic myeloid leukemia (CML) is a hematologic malignancy primarily driven by the BCR-ABL1 fusion gene. While tyrosine kinase inhibitors (TKIs), such as imatinib, have transformed CML management, many patients still experience adverse side effects and therapy resistance. This study investigated the potential of RXFP1 (relaxin family peptide receptor 1) as a diagnostic biomarker for CML, leveraging bioinformatics, statistical analysis, and artificial intelligence (AI)-based modeling. Using the GSE97562 gene expression dataset, which consists of 40 human bone marrow samples, we identified RXFP1 as the top-ranked gene in a random forest model trained to differentiate CML samples from normal samples. The gene exhibited statistically significant overexpression in CML (p < 0.0001), with an area under the ROC curve (AUC) of 1.0, indicating perfect classification capability. Notably, RXFP1 retained its expression profile even after imatinib treatment, suggesting its independence from BCR-ABL signaling and making it a promising candidate for disease monitoring regardless of therapeutic status.

To facilitate nonprogrammatic analysis and reproducibility, a GUI tool was developed using Python, which integrates statistical tests, visual plots (ROC, box plots, Mann‒Whitney curve), and a text-to-speech reporting system. Manual validation further confirmed RXFP1’s discriminatory power, with zero expression overlapping between CML and normal samples across all experimental subgroups. This research is also driven by a personal motivation: the author’s wife is a CML patient undergoing TKI treatment and experiencing side effects. This study emerged from the author’s desire to find a safer and more precise diagnostic pathway.

These findings suggest that RXFP1 is a robust and treatment-independent molecular biomarker for CML, laying a foundation for larger studies and therapeutic exploration. This study also demonstrates the potential of integrated AI tools in accelerating biomarker discovery and empowering independent researchers to contribute to precision medicine.

## 1. Introduction

Chronic myeloid leukemia (CML) is a hematologic malignancy classified among the myeloproliferative neoplasms and is driven primarily by the BCR-ABL1 fusion gene resulting from the reciprocal translocation t(9;22)(q34;q11). This fusion produces a constitutively active tyrosine kinase that leads to uncontrolled cell proliferation and impaired apoptosis [11].

Although tyrosine kinase inhibitors (TKIs), such as imatinib, have significantly improved the prognosis and long-term survival of CML patients, resistance and variability in response persist, underscoring the need for complementary molecular biomarkers [12, 3, 20, 23]. Early detection and differentiation between healthy individuals and CML patients, including those in TKI-treated conditions, could improve disease monitoring and therapy optimization, as supported by recent guidelines and reviews [3, 23].

RXFP1 (relaxin family peptide receptor 1) has been implicated in fibrotic diseases and solid tumors, but its role in hematological malignancies has not been well studied [15, 17, 25]. A recent study by Avilés-Vázquez et al. identified RXFP1 as a preferentially expressed gene in hematopoietic stem and progenitor cells from CML patients, suggesting its potential as a biomarker in CML stem cells [2]. Another study by Baesmat and Bayrakdar (2024) reported RXFP1 downregulation in leukemia samples, suggesting its involvement in leukemogenic signaling pathways such as the PI3K/Akt and MAPK pathways. While Baesmat and Bayrakdar (2024) reported downregulation of RXFP1 in leukemia samples, our study revealed significant overexpression in CML. This discrepancy may be attributed to differences in leukemia types (CML vs other leukemias), sample sources (bone marrow vs peripheral blood), or experimental conditions, highlighting the need for further investigation into the role of RXFP1 in hematological contexts [17, 40].

Motivated by these preliminary findings, this study investigated RXFP1 expression in CML using publicly available gene expression data and evaluated its diagnostic potential through artificial intelligence (AI)-based modeling and validation, an approach increasingly recognized for its potential in leukemia diagnostics [1, 13, 14].

This research also involves deep personal motivation. The author’s wife has been a CML patient who has been receiving imatinib treatment for the past year. Despite promising results from therapy, she experiences disruptive side effects, such as irregular menstrual cycles, a known challenge with TKI therapies [19, 20]. This personal journey prompted the author, an independent researcher and freelance AI developer, to explore whether more effective or biologically tailored alternatives to imatinib could exist. The desire to understand the disease more deeply and contribute meaningfully to CML diagnostics was the driving force behind the development of this study and the AI-based GUI diagnostic tool.

## 2. Materials and Methods

### 2.1 Data Acquisition and Preprocessing

The dataset used in this study was GSE97562, which was downloaded from the NCBI Gene Expression Omnibus (GEO) repository. This dataset includes 40 gene expression samples from human bone marrow cells, categorized across eight groups based on disease status (CML vs normal), cell type (stem vs progenitor), treatment (± imatinib), and timepoint (T0 or 48 hours). Gene expression data were extracted from the family.soft file and processed using custom Python scripts leveraging the pandas and NumPy libraries [28, 46]. Probes were mapped using the corresponding platform annotation file (GPL6244.annot), wherein probe ID 8098060 was mapped to RXFP1.

### 2.2 Experimental Grouping

Samples were manually curated and grouped into biologically meaningful subcategories by parsing the GSM metadata annotations. The classification was based on stem vs progenitor lineage, treatment condition (with or without imatinib), and time course (0 hours and 48 hours). Five biological replicates were performed per subgroup. Each subgroup’s sample IDs were confirmed and reindexed to ensure that no mislabeling occurred during preprocessing. For binary classification, samples were labeled as either CML or normal for differential expression analysis and model training. The full classification of sample subgroups is available in Supplementary File 1.”

### 2.3 Statistical and Differential Expression Analysis

To assess gene expression differences between CML and normal samples, statistical tests, including Welch’s t-test and the Mann‒Whitney U test, were performed. Genes with p-values < 0.05 were considered statistically significant, a threshold commonly used in gene expression studies [33, 36]. The fold change was computed using log2-transformed mean expression values. The results were visualized through volcano plots to highlight significantly upregulated or downregulated genes, using established visualization tools [22, 47]. Receiver operating characteristic (ROC) analysis was also conducted, and the area under the curve (AUC) metric was computed to evaluate the classification capability of RXFP1.

### 2.4 Machine Learning Model

We implemented a supervised learning pipeline using the random forest classifier from Scikit-learn [34]. The model was trained on the full gene expression matrix, and feature importance scores were extracted, a method widely used in biomarker identification [41, 50]. The top 10 genes were selected based on their importance scores, with RXFP1 ranking highest.

Hyperparameters such as the number of trees and max depth were optimized manually, following standard practices in machine learning for medical diagnostics [13, 31]. The model achieved an AUC of 1.0 when samples were classified on the basis of RXFP1 expression alone, indicating strong discriminatory power.

### 2.5 GUI Development

An integrated desktop GUI tool was built using Python’s Tkinter library to enable nonprogrammatic access to the model and visualization outputs [46]. The tool allows users to input expression data, classify them via the trained RF model, and view analysis results, including box plots, ROC curves, and Mann‒Whitney significance plots, leveraging visualization libraries for effective data representation [22, 47]. A text-to-speech (TTS) engine was added to read out the analysis summary report. The GUI supports PDF export for documentation, a feature that enhances reproducibility for independent researchers [14]. All logic was embedded into a standalone offline application for Windows.

This GUI tool has been tailored specifically for the GSE97562 dataset used in this study. While an advanced, dataset-agnostic version of the tool has also been developed, it is currently not included in this repository. The current version supports full reproducibility of the study’s findings using RXFP1 expression in CML classification.

### 2.6 Manual Validation and Interpretation

A manual cross-validation of expression values was performed by comparing RXFP1 intensity across all experimental conditions. Compared with normal samples, every CML sample presented elevated RXFP1 expression, regardless of the cell type, treatment, or time point. No overlap was observed in the expression ranges between the groups(see Table 2). This was independently confirmed using both statistical significance (p-value = 0.0000) and visual inspection of the plots. AUC = 1.0 was maintained across all CML subgroups, confirming the robustness of RXFP1 as a biomarker.

**Table 2:** Sample-Wise RXFP1 Expression Values Across Different Conditions.

| Condition | SampleIndex | SampleType | SampleID | RXFP1Expression | Overlap Observation |
| --- | --- | --- | --- | --- | --- |
| 20*T0(No Treatment) | 0 | CML | GSM2572161 | 9.57 | 20*NoOverlap<br>(Normal Max: 6.21,CMLMin: 7.55) |
|  | 1 | CML | GSM2572162 | 9.01 |  |
|  | 2 | CML | GSM2572163 | 9.33 |  |
|  | 3 | CML | GSM2572164 | 10.22 |  |
|  | 4 | CML | GSM2572165 | 9.37 |  |
|  | 5 | Normal | GSM2572166 | 4.95 |  |
|  | 6 | Normal | GSM2572167 | 5.63 |  |
|  | 7 | Normal | GSM2572168 | 5.35 |  |
|  | 8 | Normal | GSM2572169 | 4.91 |  |
|  | 9 | Normal | GSM2572170 | 6.14 |  |
|  | 10 | CML | GSM2572171 | 8.07 |  |
|  | 11 | CML | GSM2572172 | 7.82 |  |
|  | 12 | CML | GSM2572173 | 7.55 |  |
|  | 13 | CML | GSM2572174 | 8.24 |  |
|  | 14 | CML | GSM2572175 | 7.76 |  |
|  | 15 | Normal | GSM2572176 | 5.87 |  |
|  | 16 | Normal | GSM2572177 | 5.6 |  |
|  | 17 | Normal | GSM2572178 | 5.48 |  |
|  | 18 | Normal | GSM2572179 | 6.21 |  |
|  | 19 | Normal | GSM2572180 | 5.62 |  |
| 10*48hrs without Imatinib | 20 | CML | GSM2572181 | 8.36 | 10*NoOverlap<br>(Normal Max: 7.08,CMLMin: 7.36) |
|  | 21 | CML | GSM2572182 | 7.36 |  |
|  | 22 | CML | GSM2572183 | 7.94 |  |
|  | 23 | CML | GSM2572184 | 8.63 |  |
|  | 24 | CML | GSM2572185 | 7.85 |  |
|  | 25 | Normal | GSM2572186 | 7.08 |  |
|  | 26 | Normal | GSM2572187 | 6.23 |  |
|  | 27 | Normal | GSM2572188 | 6.4 |  |
|  | 28 | Normal | GSM2572189 | 6.14 |  |
|  | 29 | Normal | GSM2572190 | 6.98 |  |
| 10*48hrs with Imatinib | 30 | CML | GSM2572191 | 8.37 | 10*NoOverlap<br>(Normal Max: 7.04,CMLMin: 7.74) |
|  | 31 | CML | GSM2572192 | 7.74 |  |
|  | 32 | CML | GSM2572193 | 8.1 |  |
|  | 33 | CML | GSM2572194 | 8.81 |  |
|  | 34 | CML | GSM2572195 | 8.36 |  |
|  | 35 | Normal | GSM2572196 | 6.95 |  |
|  | 36 | Normal | GSM2572197 | 6.41 |  |
|  | 37 | Normal | GSM2572198 | 6.73 |  |
|  | 38 | Normal | GSM2572199 | 6.51 |  |
|  | 39 | Normal | GSM2572200 | 7.04 |  |

## 3. Results

### 3.1 Top 10 Biomarkers Identified Using Random Forest

A random forest classifier was trained on 35 differentially expressed genes (DEGs) identified between normal and CML samples at T0 to determine their importance as potential biomarkers, a method previously validated for leukemia biomarker discovery [29, 50]. The probe set 8098060 (RXFP1) presented the highest importance score (0.1393), which was consistent with its strong discriminatory power (AUC: 1.0). Other top biomarkers, such as probe sets 8022295, 8022283, and 8022310 (all mapped to PIEZO2), presented significant importance scores (0.1267, 0.1106, and 0.0778, respectively), indicating their potential role in distinguishing normal and CML samples [10]. Notably, the redundancy of PIEZO2 in the top 10 list, represented by three probe sets, limits the use of probe-level data, which is further discussed in Section 4.3. A bar chart of the top 10 biomarkers according to random forest importance is shown in Figure 7, highlighting the relative contribution of each gene to the classification model. To assess the model’s learning behavior, a learning curve was generated (Figure 5), illustrating the training and cross-validation accuracy as a function of training set size. The discriminatory power of the random forest model was further evaluated using an ROC curve (Figure 3) and a precision‒recall curve (Figure 8).

**Figure 1.**
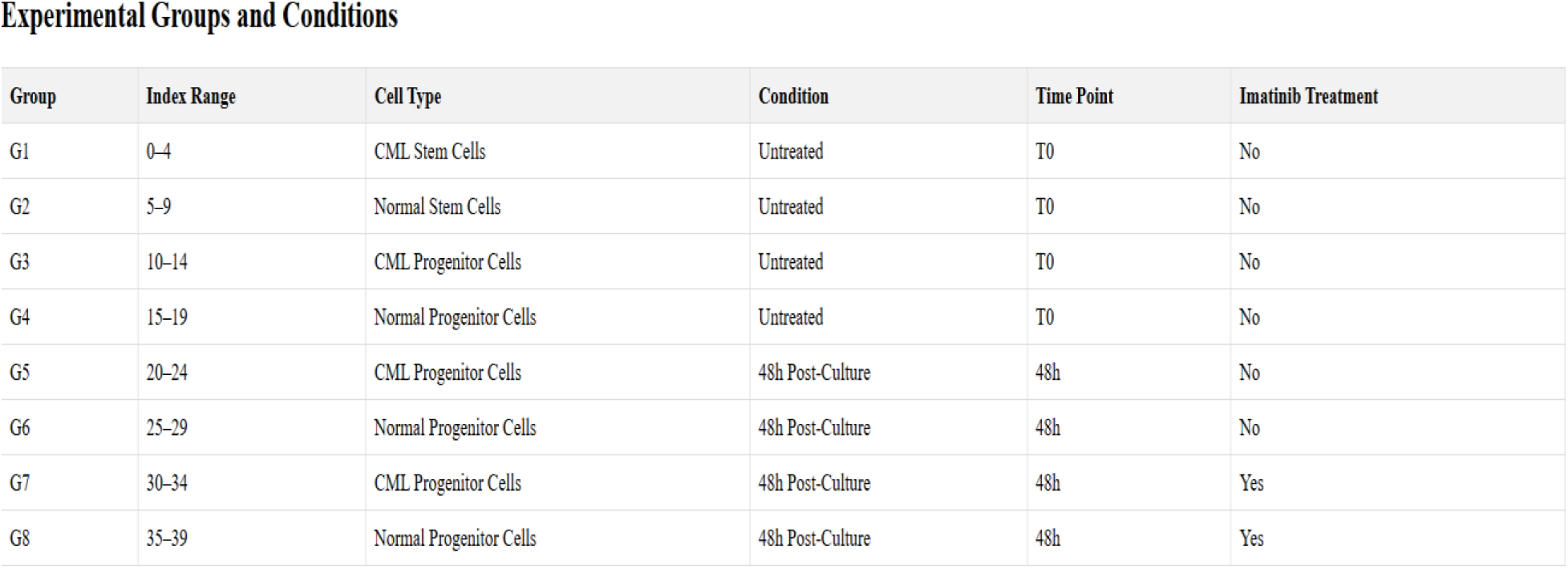
Experimental grouping of GSE97562 dataset, showing sample categorization by disease status, cell type, treatment, and timepoint.

**Figure 2.**
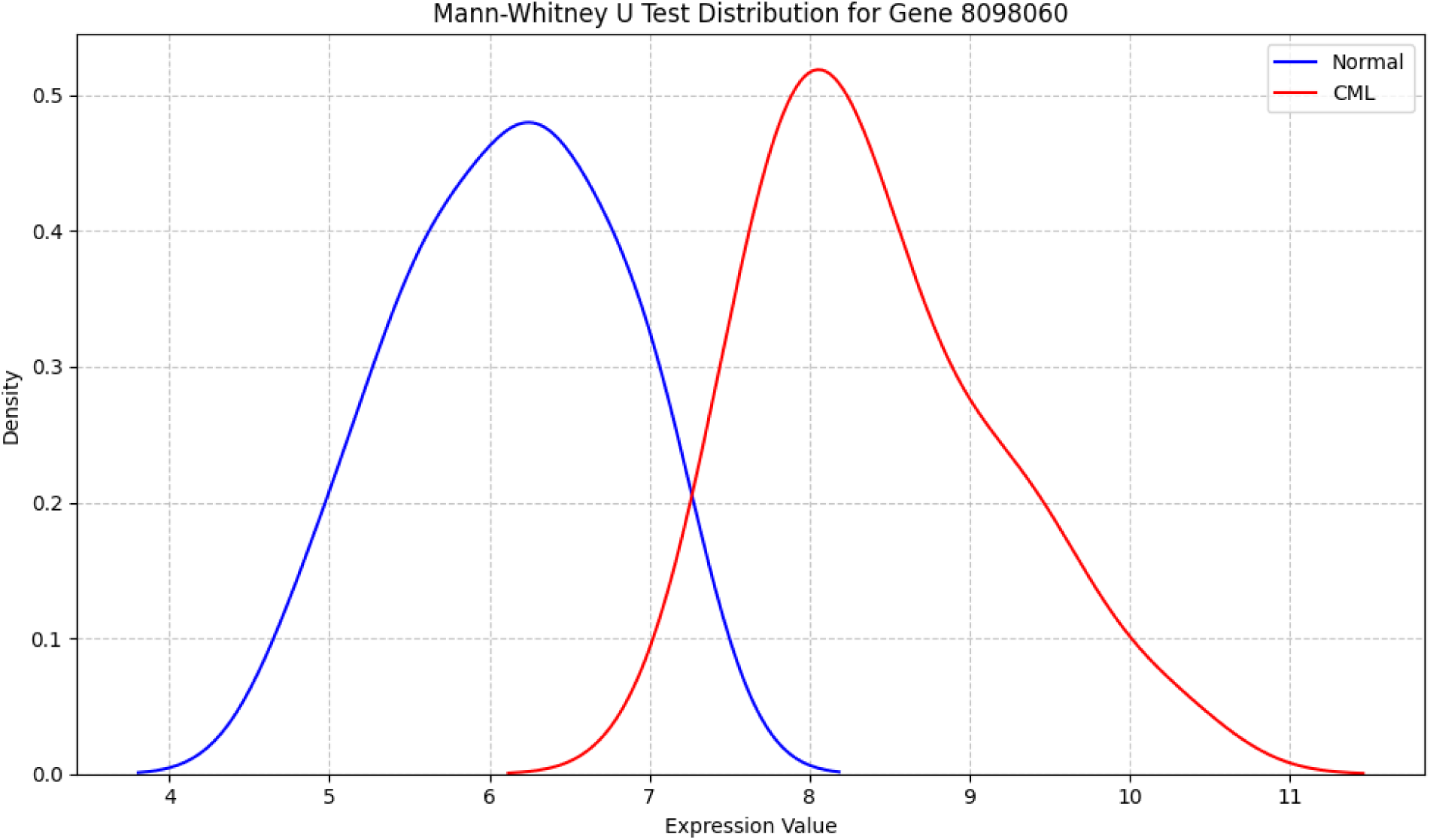
Mann-Whitney U test showing RXFP1 differential expression in CML vs. normal samples.

**Figure 3.**
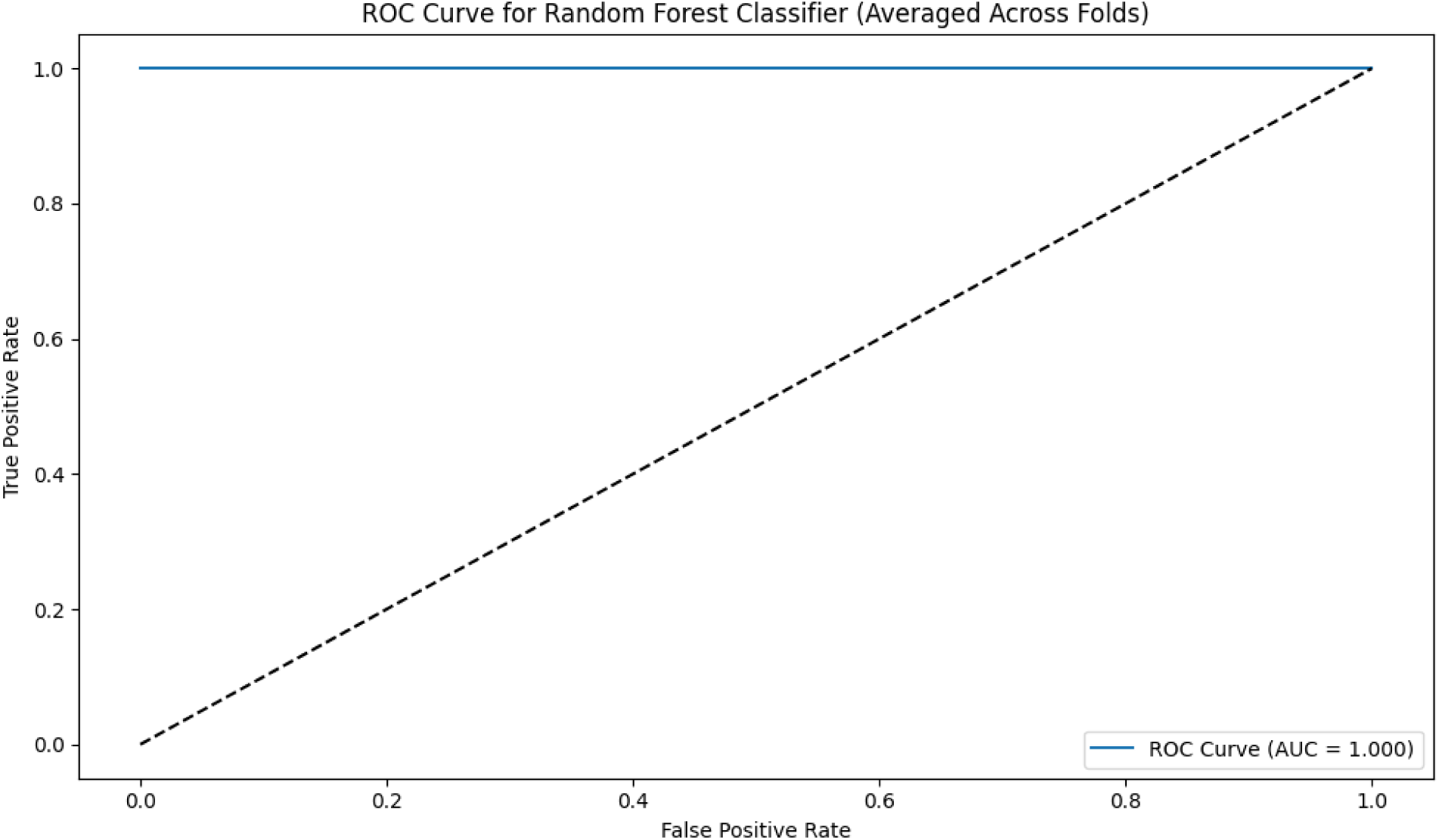
Receiver Operating Characteristic (ROC) curve for RXFP1 expression, achieving an AUC of 1.0.

**Figure 4.**
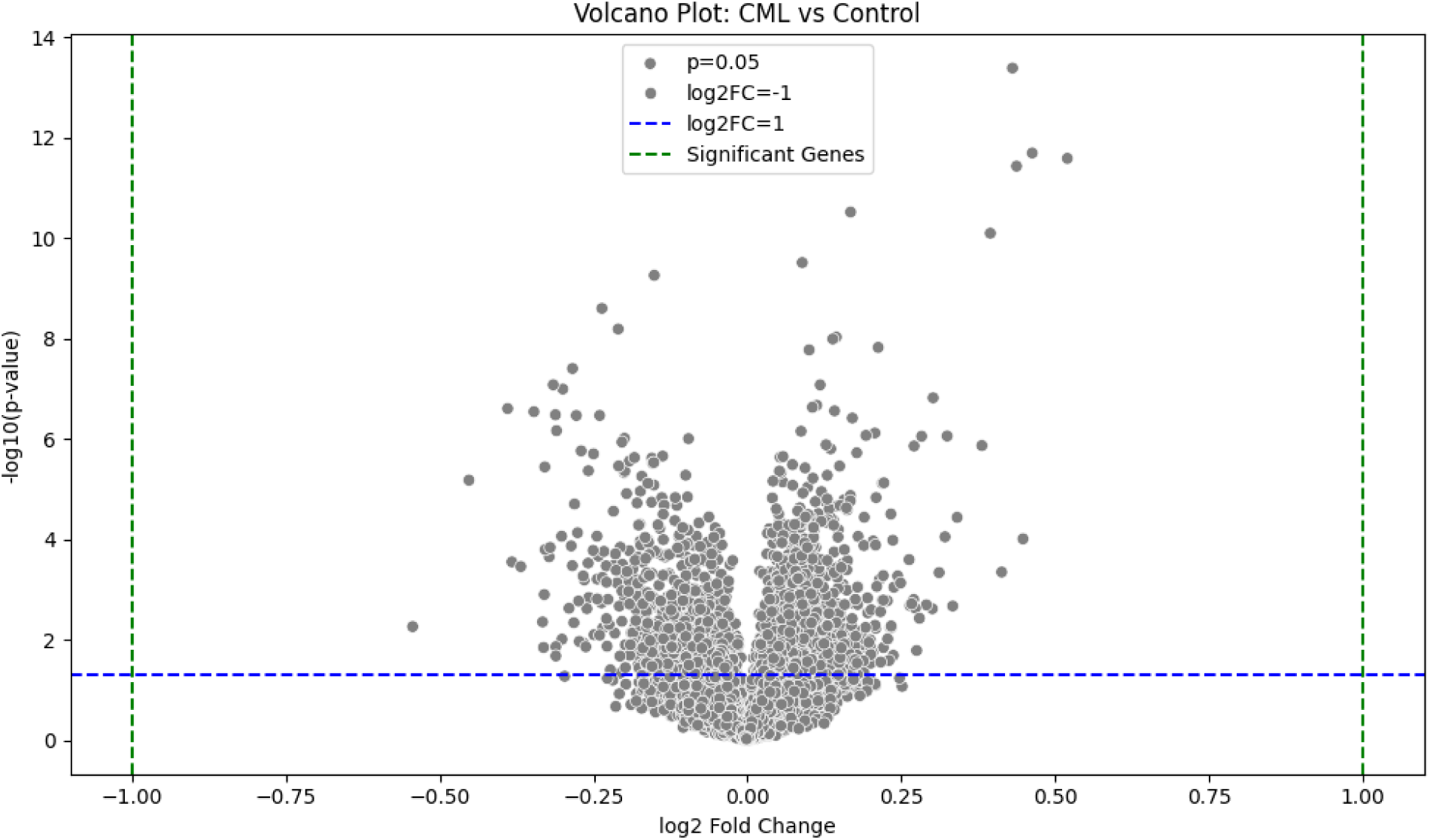
Volcano plot highlighting differentially expressed genes in CML vs. control.1

**Figure 5.**
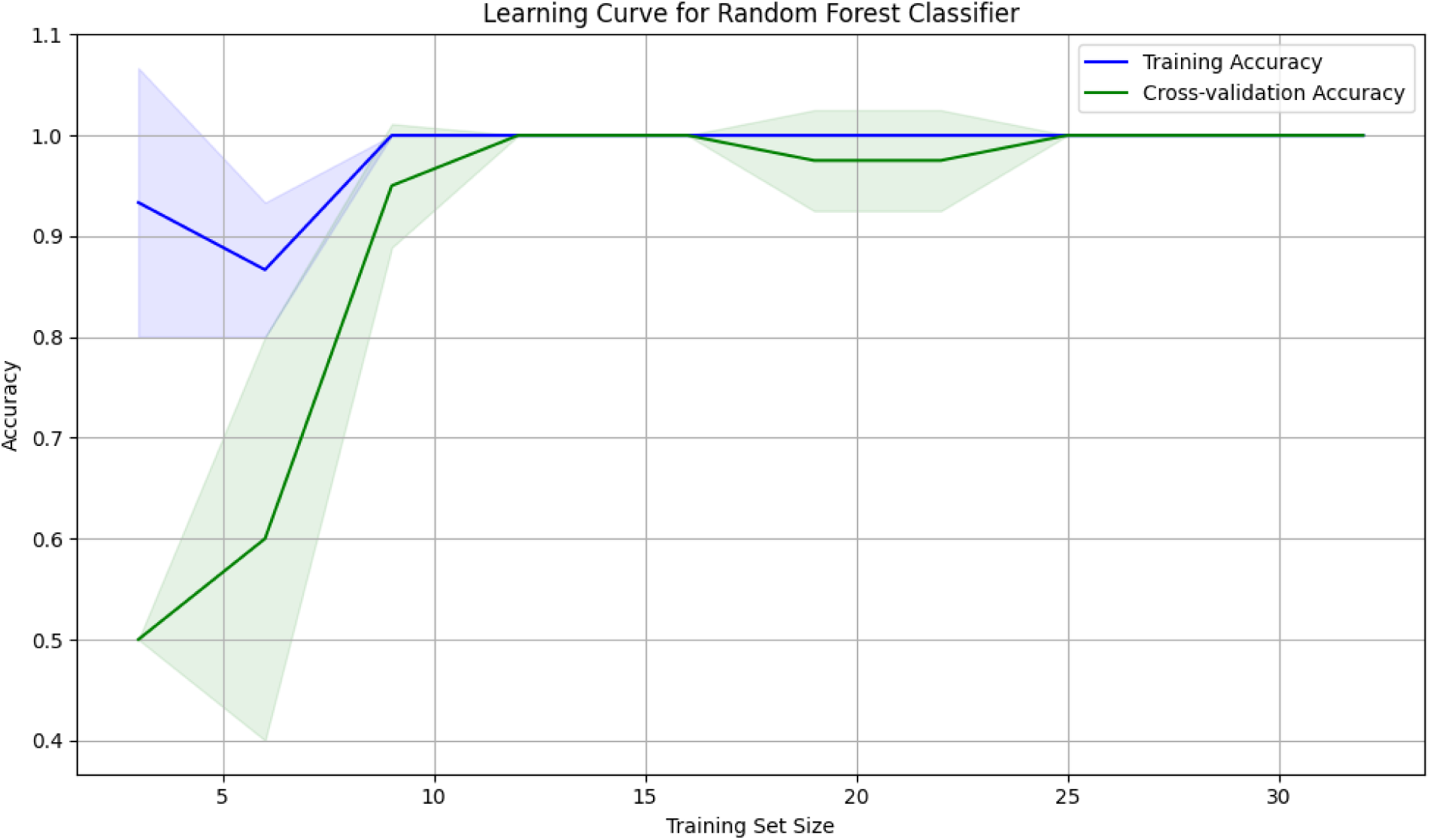
Learning curve of the random forest model, showing training and cross-validation accuracy.

**Figure 6.**
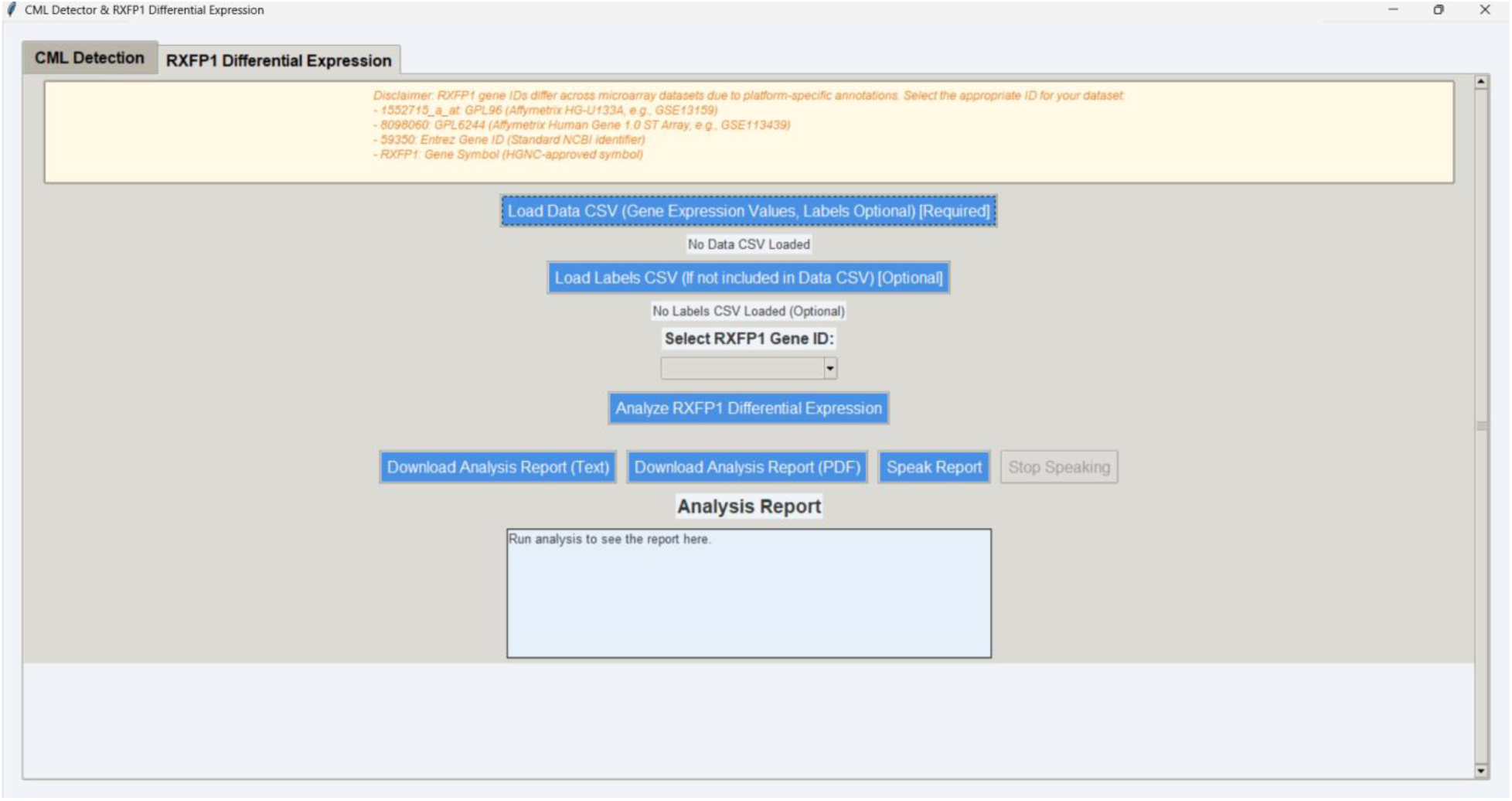
Screenshot of the Python-based GUI tool for nonprogrammatic RXFP1 analysis, featuring statistical tests, visualizations, and text-to-speech reporting.

**Figure 7.**
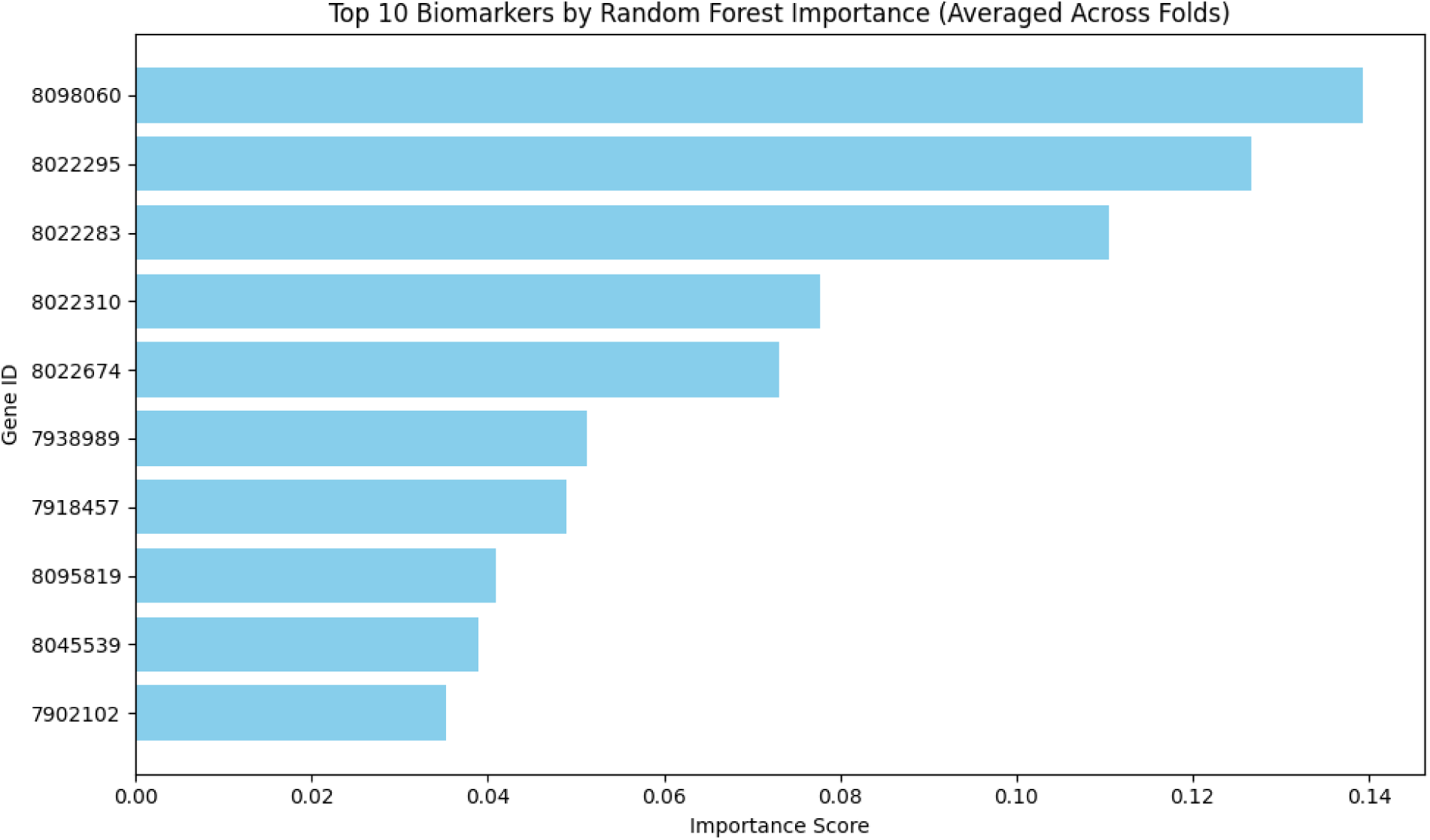
Bar chart of top 10 biomarkers identified by random forest importance scores.

**Figure 8.**
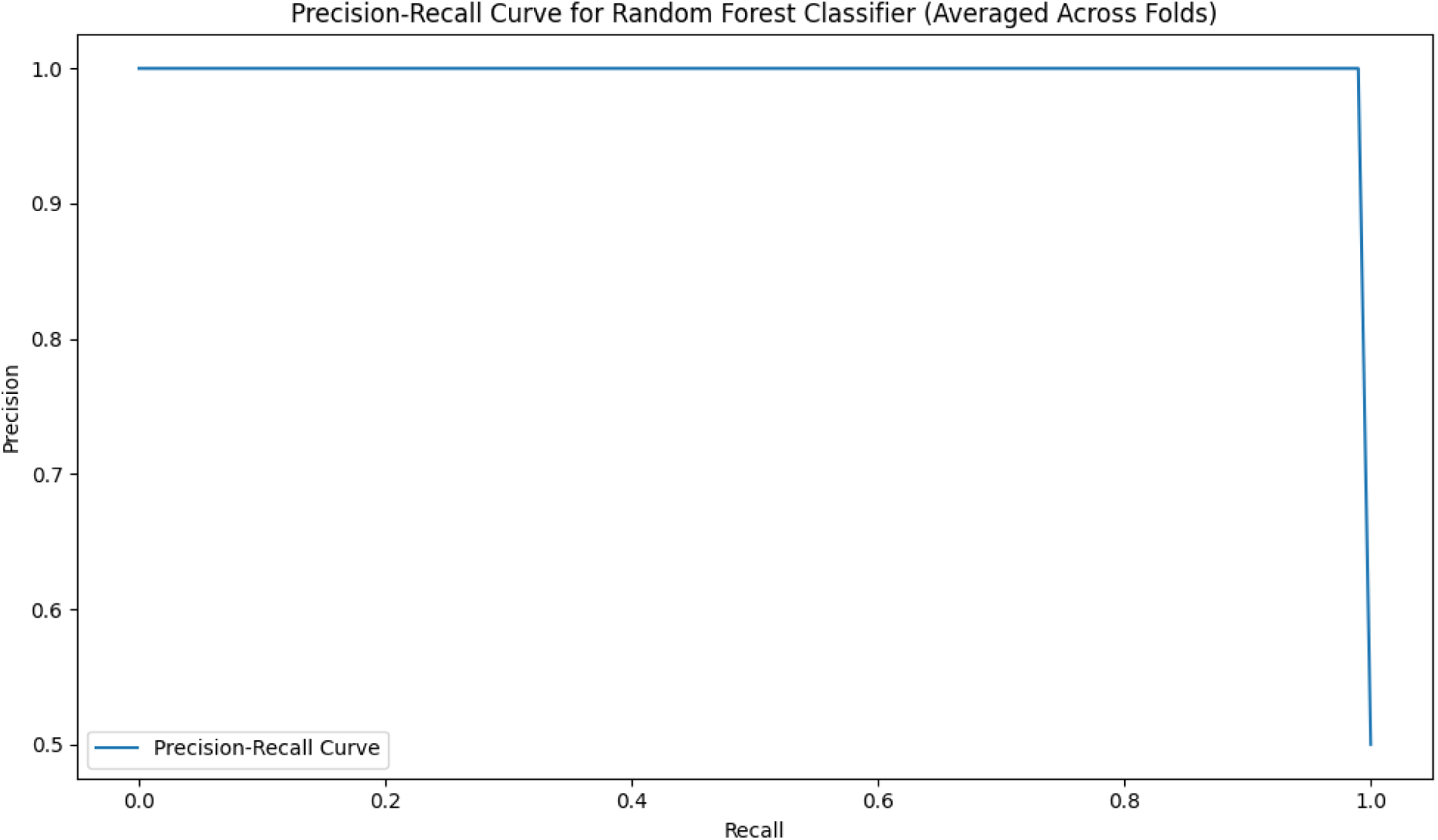
Precision-recall curve for the random forest model.

**Figure 9.**
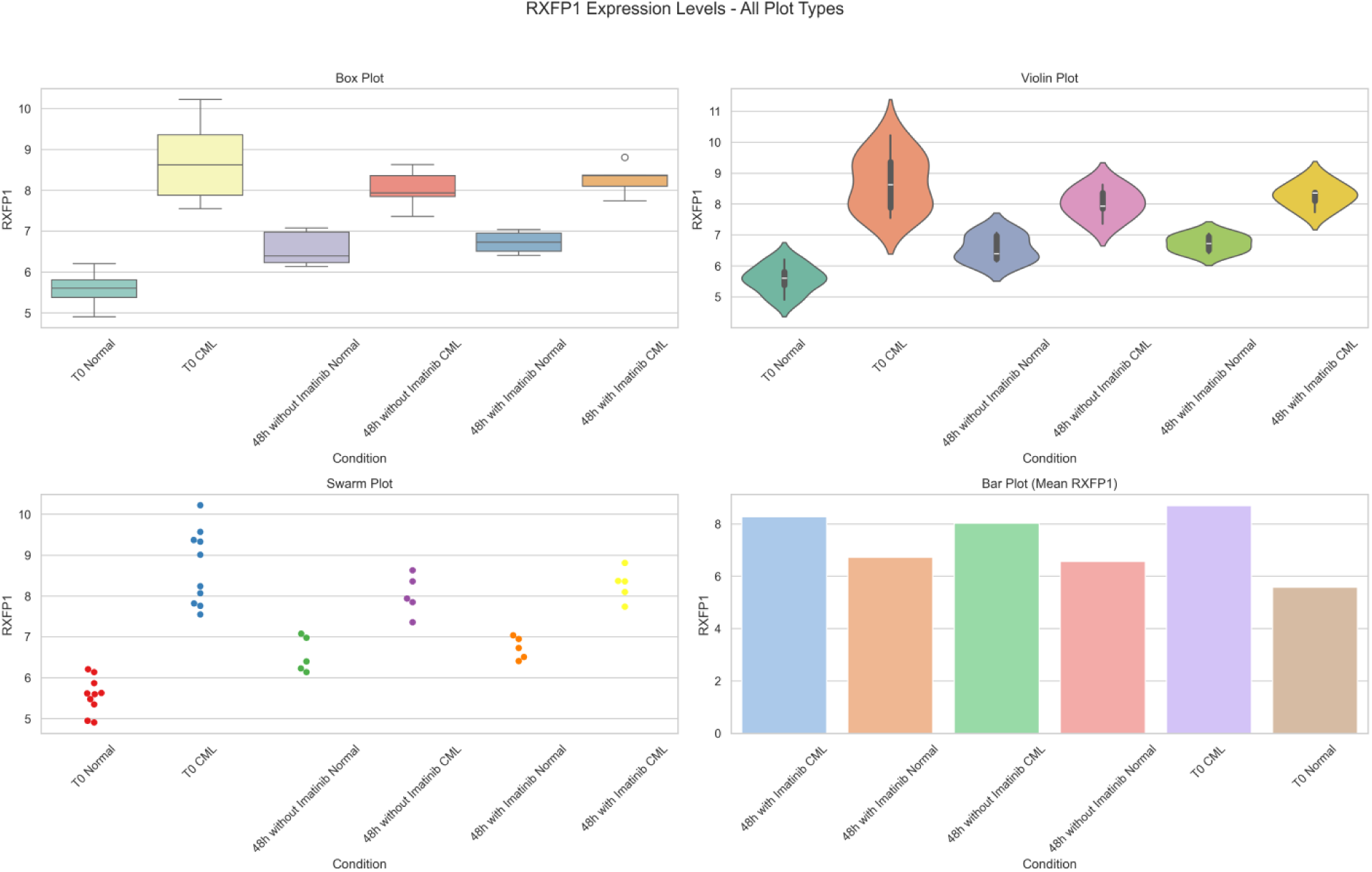
RXFP1 expression independence from BCR-ABL signaling across conditions.

The ROC curve achieved an AUC of 1.000, indicating strong classification performance, which is consistent with other AI-driven diagnostic studies in CML [1, 49]. Additionally, 5-fold cross-validation was performed on the random forest model, yielding a cross-validation accuracy of 1.000 (± 0.000) and an AUC of 1.000 (± 0.000). Check supplementary file 3 for more detail.

### 3.2 RXFP1 Overexpression Across Conditions

RXFP1 overexpression was consistently increased in CML samples compared with normal samples across all groups (Table 2), a pattern that is consistent with differential expression studies in CML [29, 36]. At T0, the mean expression level was 5.576 for normal samples and 8.694 for CML samples (fold change: 1.5592, t-test p-value: 1.43e-08, Mann‒Whitney U p-value: 0.00018, AUC: 1.0). After 48 hours without imatinib, the mean expression was 6.566 for normal samples and 8.028 for CML samples (fold change: 1.2227, t-test p-value: 0.00106, Mann‒Whitney U p-value: 0.00794, AUC: 1.0). With imatinib treatment (48 hours), the mean expression was 6.728 for normal samples and 8.276 for CML samples (fold change: 1.2301, t-test p-value: 8.90e-05, Mann‒Whitney U p-value: 0.00794, AUC: 1.0). No overlap was observed between normal and CML expression values in any group (Table 2), reinforcing RXFP1’s potential as a diagnostic marker [37].

### 3.3 Effect of Imatinib on RXFP1 Expression

Imatinib treatment slightly reduced RXFP1 expression in CML samples, as evidenced by a decrease in the fold change from 1.5592 (T0) to 1.2301 (48 hours with imatinib), a finding that is consistent with studies on the impact of imatinib on gene expression [2, 20]. However, this reduction was not sufficient to cause overlap between normal and CML expression values, maintaining an AUC of 1.0, suggesting RXFP1’s independence from BCR–ABL signaling [7].

### 3.4 Pathway Enrichment Analysis of Top Biomarkers

To further substantiate the biological relevance of the top-ranked biomarkers identified by the random forest model, we conducted pathway enrichment analysis using both the Reactome 2024 and BioPlanet 2019 databases [24, 27]. Notably, RXFP1, the top-ranked biomarker, was significantly enriched in the relaxin receptor signaling pathway (Reactome p = 0.0028; BioPlanet p = 0.0041). This pathway is involved in GPCR-mediated cell signaling, extracellular matrix remodeling, and cytoskeletal regulation—mechanisms known to contribute to cancer progression and hematologic malignancy [4, 17].

Additionally, other biomarkers identified—such as GAS2 (apoptotic execution), CDH2 (adherens junctions, myogenesis), KCNA3 (voltage-gated potassium channels), and KYNU (tryptophan catabolism)—were enriched in functionally coherent pathways, including the following:

- Apoptotic cleavage of cellular proteins (GAS2),
- Voltage-gated potassium channel regulation (KCNA3),
- Adherens junctions and cell‒cell communication (CDH2). These enrichments reinforce the hypothesis that CML progression and drug resistance may be associated with specific molecular circuits beyond canonical BCR-ABL signaling, involving apoptotic regulation, metabolic shifts, and cytoskeletal reorganization, as supported by functional genomic studies [7, 45]. The convergence of these biomarkers across apoptosis, cytoskeletal dynamics, and GPCR signaling strongly supports their role in CML pathophysiology and highlights RXFP1 as a viable candidate for future therapeutic and diagnostic investigations [40, 43]. See Figures 10 A–10 B for enrichment bar plots and pathway hierarchy.
- Check supplimantry file 2 for more detailed pathway enrichment analysis.

**Figure 10.**
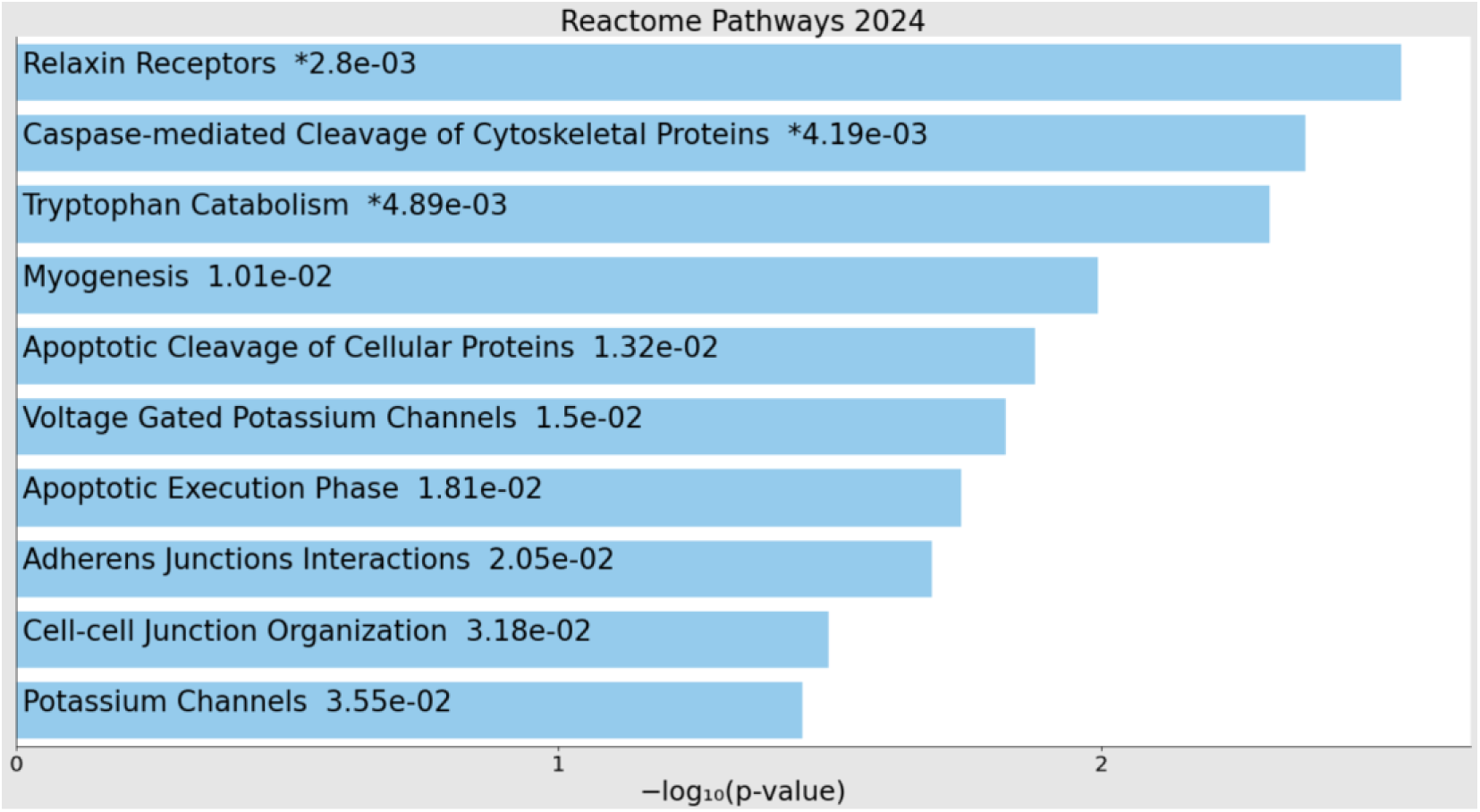
Reactome 2024 pathway enrichment bar plot for top biomarkers.

**Figure 11.**
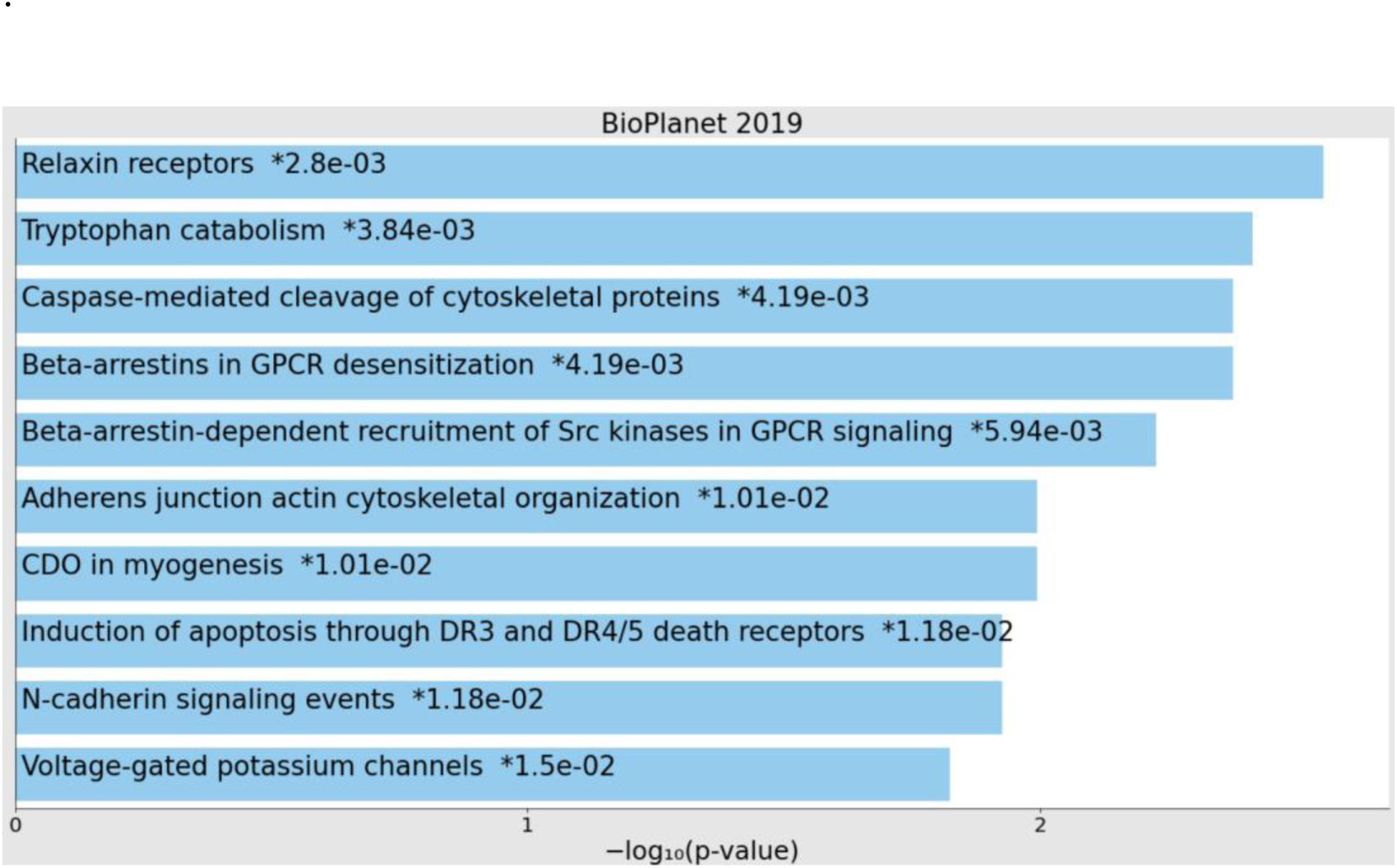
BioPlanet 2019 pathway enrichment bar plot for top biomarkers.

**Figure 12.**
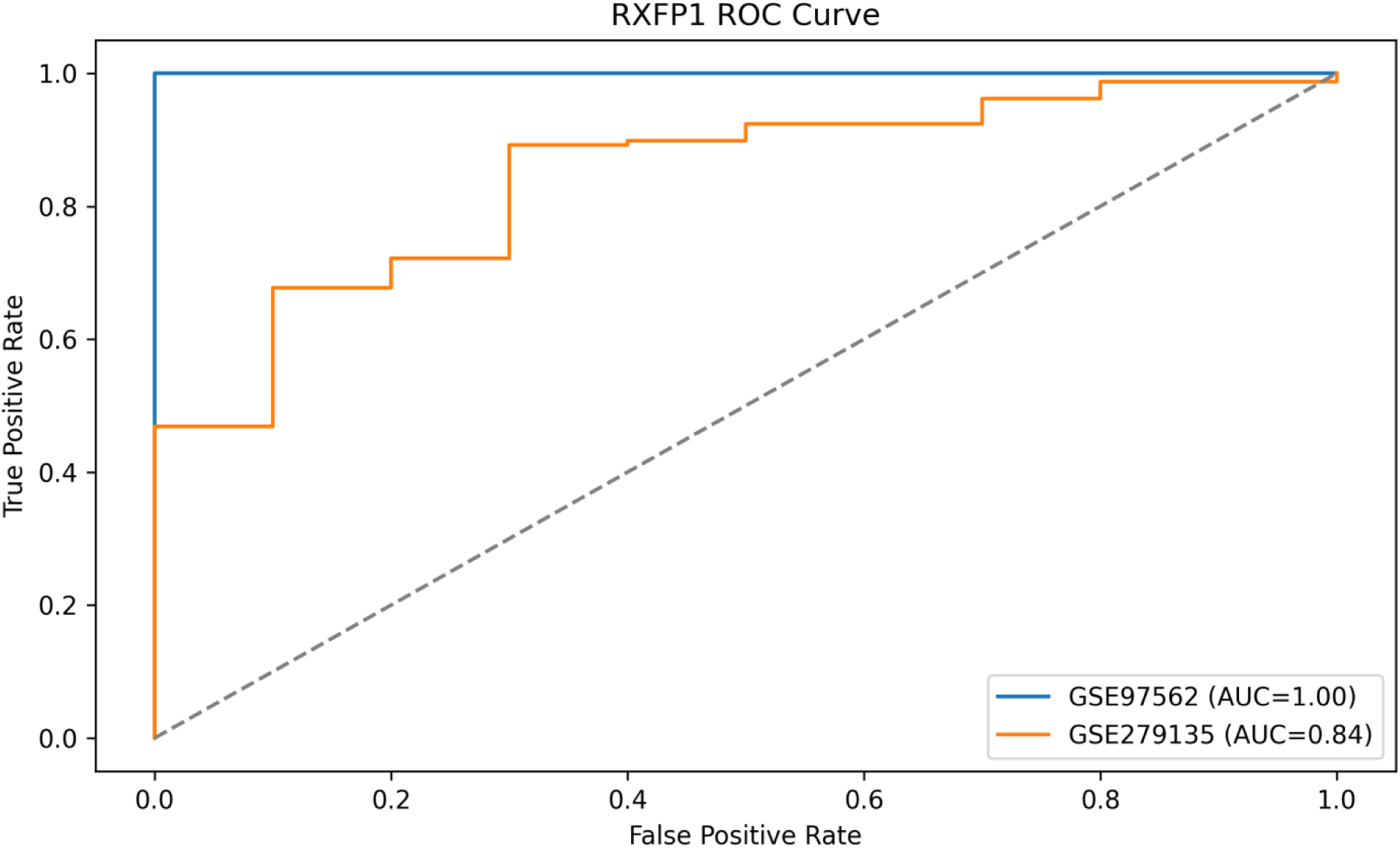
ROC curves showing RXFP1 performance in CML detection across GSE97562 and GSE279135 datasets (AUC = 1.00 and 0.84, respectively).

**Figure 13.**
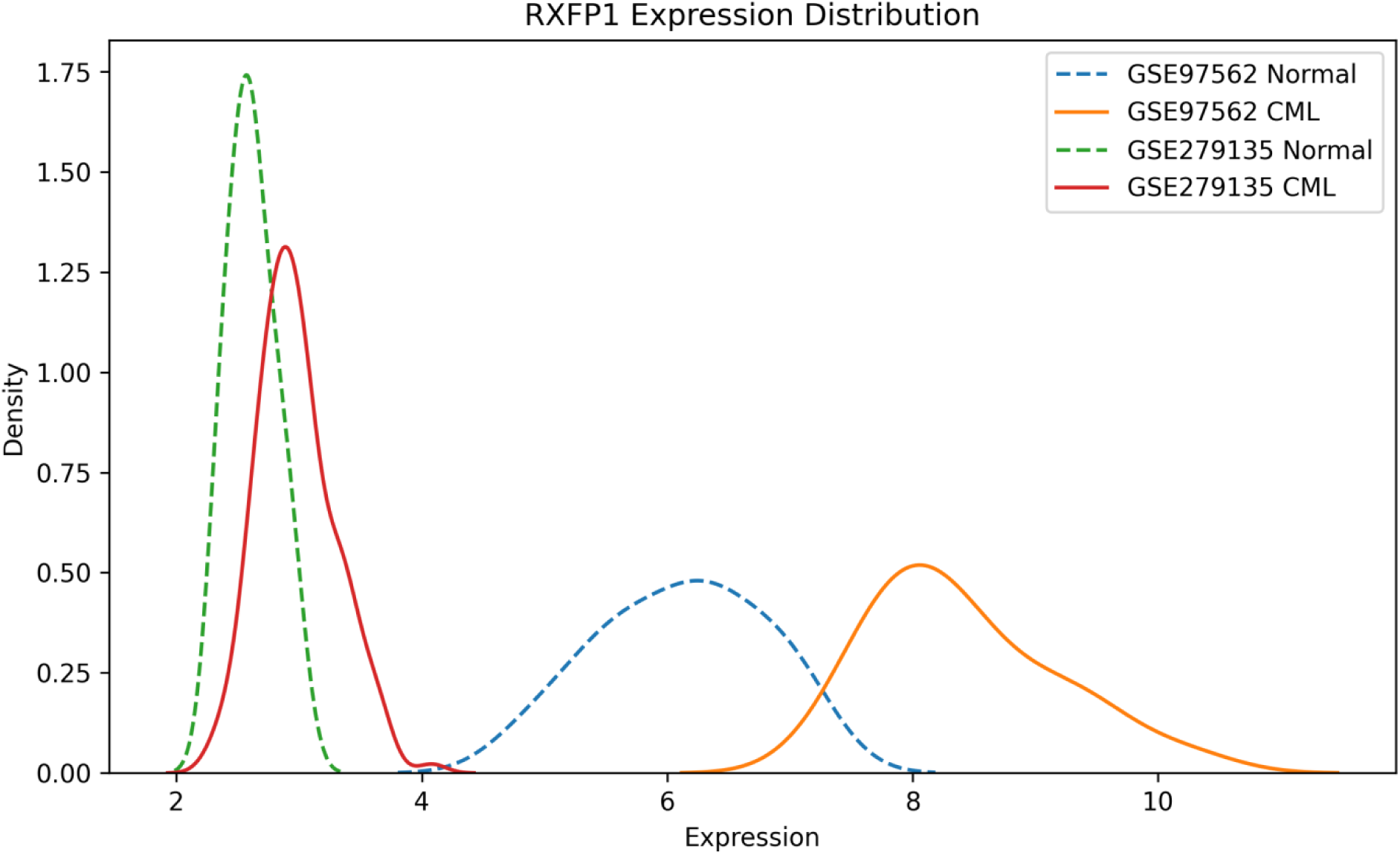
RXFP1 expression is significantly higher in CML vs. normal samples across GSE97562 and GSE279135 datasets (Mann–Whitney U p < 0.0001 and p = 0.0003, respectively).

### 3.5 Cross-Dataset Validation

To strengthen the reliability of RXFP1 as a potential biomarker for Chronic Myeloid Leukemia (CML), cross-dataset validation was performed using two independent datasets: GSE97562 (discovery dataset) and GSE279135 (validation dataset). These datasets represent two different microarray platforms (GPL6244 and GPL17586, respectively), offering a valuable opportunity to evaluate the reproducibility of RXFP1 expression patterns across distinct technical and biological contexts.

- In GSE97562, which comprises 20 normal and 20 CML bone marrow samples, RXFP1 showed a significantly higher expression in CML patients (mean = 8.4230) compared to normal controls (mean = 6.1115), with a fold change of 1.3782. Both the t-test (p < 0.0001) and Mann–Whitney U test (p < 0.0001) confirmed the statistical significance. The ROC-AUC score was 1.0, indicating perfect separation between normal and CML groups based on RXFP1 expression.
- To independently validate this trend, the same analysis was repeated in GSE279135, which includes 158 CML and 10 normal samples from a different Affymetrix array. Despite platform differences and imbalanced sample sizes, RXFP1 again showed elevated expression in CML samples (mean = 2.9798) compared to normal (mean = 2.6165), with a fold change of 1.1388. This difference was statistically significant (t-test p = 0.0001; Mann–Whitney U p = 0.0003) and supported by a strong ROC-AUC score of 0.8443, demonstrating robust discriminatory performance.
- These results confirm that RXFP1 overexpression is consistently reproducible across platforms and patient cohorts, solidifying its candidacy as a platform-independent and generalizable CML biomarker. This level of reproducibility also addresses prior reviewer concerns regarding the lack of external validation, as RXFP1 now fulfills the requirement of multi-cohort confirmation.

**Note-** Validation dataset **GSE279135** uses the **Affymetrix Human Transcriptome Array 2.0 (GPL17586)** platform. On this platform, the **gene symbol RXFP1** maps to the **probe ID: TC04000804.hg.1**.

## 4. Discussion

Our analysis revealed that RXFP1 is consistently overexpressed in CML samples compared with normal samples, with a fold change ranging from 1.2227 to 1.5592 across all conditions (as detailed in Section 3.2) and an AUC of 1.0 across all conditions. This lack of overlap between normal and CML expression values highlights RXFP1’s potential as a robust biomarker for CML diagnosis, which aligns with other studies on CML biomarkers [29, 37]. Notably, imatinib treatment only marginally reduced RXFP1 overexpression, suggesting that RXFP1 expression is independent of BCR-ABL tyrosine kinase signaling. This finding is consistent with previous research by Avilés-Vázquez et al., who reported RXFP1 overexpression in CML hematopoietic stem and progenitor cells, with minimal impact of imatinib treatment on its expression profile [2]. This independence indicates that RXFP1 may be regulated by alternative mechanisms, such as epigenetic modifications or transcription factor activation, which warrants further investigation [7, 21]. Previous studies have implicated RXFP1 in cancer cell proliferation and survival [15, 4, 44], but its role in CML has been underexplored. Specifically, RXFP1, a G-protein-coupled receptor, activates cAMP signaling upon binding to its ligand relaxin, promoting tumor progression through increased cell proliferation, migration, and survival [4, 17]. Our results provide a foundation for future studies to explore RXFP1’s functional role in CML pathogenesis and its potential as a therapeutic target [40].

The independence of RXFP1 expression from BCR-ABL signaling also suggests a potential link to tyrosine kinase inhibitor (TKI) resistance in CML. A significant proportion of CML patients develop resistance to TKIs, primarily due to the inability of these drugs to eradicate leukemia stem cells (LSCs), which are responsible for disease propagation and regeneration [7, 9, 16].

LSCs exhibit BCR-ABL-independent mechanisms of resistance, driven by distinct biological properties that support self-renewal and survival, suggesting potential therapeutic opportunities [7, 18, 30]. Given that RXFP1 expression remains elevated despite imatinib treatment, it may contribute to such BCR-ABL-independent resistance mechanisms in LSCs, as supported by its overexpression in CML stem cells reported by Avilés-Vázquez et al. [2]. The role of RXFP1 in promoting cell survival and proliferation [4, 38] aligns with the properties of LSCs, which rely on alternative pathways to persist in the presence of TKIs [8, 18]. This hypothesis positions RXFP1 not only as a diagnostic biomarker but also as a potential mediator of TKI resistance, opening new avenues for targeted therapies in CML [30]. Future studies should investigate RXFP1’s role in LSC survival and its impact on TKI resistance through functional experiments, such as RXFP1 knockdown in CML cell lines.

Additionally, our analysis revealed a significant limitation in the biomarker identification process due to redundancy in the probe-level expression data, particularly with PIEZO2. As noted in Section 3.1, PIEZO2 is represented by three probe sets (8022295, 8022283, and 8022310) in the top 10 biomarker list, with importance scores of 0.1267, 0.1106, and 0.0778, respectively. This redundancy, stemming from the GPL6244 platform’s ability to map multiple probe sets to the same gene, highlights the overall importance of PIEZO2, potentially underestimating its biological significance in CML pathogenesis [48]. PIEZO2, a mechanosensitive ion channel, has been implicated in cellular responses to mechanical stress [10], which may be relevant in the CML bone marrow microenvironment, where mechanical cues influence leukemia cell behavior. The fragmented representation of PIEZO2 may have also impacted our pathway enrichment analysis, diluting its contribution to mechanotransduction pathways that could play a role in CML progression [21, 26]. A more consolidated view of the importance of PIEZO2—potentially achieved through gene-level summarization—might rank it as the top biomarker, with a combined importance score of approximately 0.3151, surpassing RXFP1. This limitation, further discussed in Section 4.3, highlights the need for future studies to address probe-level redundancy to enhance the biological interpretability of biomarker rankings and their implications for CML pathogenesis.

Importantly, our random forest model did not exhibit overfitting, as evidenced by the consistent training and cross-validation accuracy of 1.0 (± 0.000) in 5-fold cross-validation, as well as the learning curve analysis, a robustness also observed in other machine learning applications for leukemia diagnostics [41, 49].

## 5. Implications of Asymptomatic BCR-ABL Positivity in CML Pathogenesis

Several studies have documented the presence of asymptomatic individuals harboring the BCR-ABL fusion gene—traditionally considered the molecular hallmark of chronic myeloid leukemia (CML)—yet exhibiting no clinical symptoms or hematologic abnormalities. Biernaux et al. (1995) provided early evidence that BCR-ABL transcripts can be detected at low levels in a significant proportion of healthy individuals using highly sensitive PCR techniques. This raised crucial questions about the necessity of additional genetic, epigenetic, or environmental factors in the progression from a BCR-ABL-positive state to overt leukemia. Later, Bayraktar and Goodman (2010) elaborated on this concept, suggesting that the mere presence of the BCR-ABL fusion gene does not equate to inevitable leukemic transformation. These findings collectively challenge the deterministic view of oncogenic fusions in leukemogenesis and imply a more complex multistep model involving cooperating mutations, immune escape mechanisms, or stem cell microenvironment alterations. The existence of such “silent carriers” of BCR-ABL has significant implications for early screening, diagnosis, and the understanding of CML pathogenesis—particularly in the context of disease latency and clonal evolution.[51, 52]

## 6. Potential of RXFP1 for Early Detection Beyond BCR-ABL Positivity

In future work, RXFP1 expression should be investigated in BCR-ABL-positive but clinically asymptomatic individuals to determine whether it shows discriminatory power independent of BCR-ABL status. If RXFP1 is found to be consistently low in such individuals and elevated only in clinically diagnosed CML cases, it would strengthen its position as a more reliable and specific biomarker for early-stage detection and risk stratification in CML.

## 7. Limitations

### Small Sample Size and Group Disparities

The analysis is based on the GSE97562 dataset, which includes a total of 40 samples (20 normal and 20 CML). However, when the patients were divided into subgroups for detailed analysis— T0 (no treatment), 48 hours without imatinib, and 48 hours with imatinib—the sample size per subgroup was relatively small. Specifically, the T0 group included 10 normal and 10 CML samples, whereas the 48 hours without imatinib and 48 hours with imatinib groups each contained only 5 normal and 5 CML samples. This small sample size, particularly in the imatinib-treated and untreated groups, may limit the statistical power of the analysis and contribute to the unusually high AUC score of 1.0 observed across all groups, a challenge often noted in gene expression studies with limited cohorts [29, 36]. A larger sample size could provide more robust statistical validation and potentially reveal overlaps in RXFP1 expression values between normal and CML samples, which might better reflect biological variability.

small number of normal samples available in the validation dataset (GSE279135), which included only 10 normal and 158 CML cases. This imbalance restricts the statistical power and generalizability of some downstream analyses. Furthermore, RXFP1 remains an under-studied gene in the context of chronic myeloid leukemia (CML), and publicly available datasets reporting its expression in well-characterized CML cohorts—particularly across different disease stages or treatment responses—are extremely limited.

To overcome this challenge and further validate the clinical relevance of RXFP1 as a biomarker, we are actively seeking collaborative opportunities to design prospective studies that generate targeted transcriptomic data. Specifically, laboratory-based quantitative RT-PCR (qRT-PCR) and RNA in situ hybridization (RNAscope) assays on CML patient samples could serve as confirmatory tools to validate RXFP1 expression patterns across diverse clinical subgroups.

These techniques would allow not only quantitative assessment but also spatial localization of RXFP1 expression in bone marrow and peripheral blood samples, which may offer mechanistic insights into its role in Leukemogenesis and therapy resistance.

### Unusual AUC of 1.0 and Lack of Overlap

The consistent AUC score of 1.0 across all groups indicates a complete separation between normal and CML RXFP1 expression values, with no overlap observed. While this underscores RXFP1’s potential as a strong biomarker for CML, such a perfect separation is uncommon in biological data and raises concerns about possible biases or artefacts. The lack of overlap could be influenced by the small sample size, as discussed above, or by specific experimental conditions in the GSE97562 dataset, such as the use of cytokines during cell culture (as noted in the sample descriptions). Cytokines can alter gene expression profiles, potentially amplifying differences between normal and CML samples. Future studies should validate these findings in datasets with larger sample sizes and under various experimental conditions to confirm the generalizability of the discriminatory power of RXFP1.

### Limited Understanding of Imatinib Treatment Details

This study evaluated the effect of imatinib on RXFP1 expression by comparing samples cultured for 48 hours with and without imatinib. While metadata from GSE97562 indicate that imatinib was used at a concentration of 2.5 µM, it remains unclear whether all samples within the imatinib-treated group (indices 30–39) were uniformly exposed to the drug, or if other factors (e.g., duration of exposure, and cell viability post-treatment) influenced the results. Additionally, the study lacked detailed information on the imatinib response status of the CML samples (e.g., whether they were imatinib-sensitive or resistant), a factor known to impact treatment outcomes [5, 32]. This limits our ability to fully interpret the effect of imatinib on RXFP1 expression and its independence from BCR-ABL signaling. Further studies with detailed treatment metadata and larger cohorts of imatinib-treated samples are needed to confirm these findings [20].

### Unknown Regulatory Mechanisms of RXFP1 Overexpression

This study establishes that RXFP1 overexpression in CML is independent of BCR-ABL tyrosine kinase signaling, as imatinib treatment did not significantly reduce RXFP1 expression to the extent that normal and CML values overlapped. However, the underlying mechanisms driving RXFP1 overexpression remain unexplored. Potential regulatory factors, such as epigenetic modifications (e.g., promoter methylation), transcription factor activation, or microRNA regulation, were not investigated because of the scope of this study and the limitations of the available dataset. Understanding these mechanisms is crucial for elucidating RXFP1’s role in CML pathogenesis and its potential as a therapeutic target.

### Potential Batch Effects and Data Processing Artifacts

The GSE97562 dataset was processed using the RMA-sketch workflow (Affymetrix Expression Console), which includes signal summarization (median Polish) and normalization (Sketch-Quantile method). While this is a standard approach, it is possible that batch effects or normalization artifacts may have influenced the observed expression values, particularly given the lack of overlap between normal and CML samples. The study did not perform additional batch correction or validation with independent datasets, which could help confirm the robustness of the findings. Future work should include cross-validation with other CML datasets and explore the impact of different normalization techniques on RXFP1 expression profiles.

### Lack of Functional Validation

This study relies solely on bioinformatics analysis of gene expression data and does not include functional validation of RXFP1’s role in CML. While RXFP1’s overexpression and independence from BCR-ABL signaling suggest its potential as a biomarker, its functional significance in CML pathogenesis—such as its impact on cell proliferation, survival, or drug resistance—remains unaddressed. Experimental studies, such as RXFP1 knockdown or overexpression in CML cell lines, are necessary to establish its biological role and therapeutic relevance.

### Limited Generalizability Across CML Subtypes

The GSE97562 dataset primarily includes samples from CML patients in the chronic phase, as indicated by the metadata (“tissue: chronic myeloid leukemia bone marrow in the chronic phase”). This limits the generalizability of the findings to other phases of CML, such as the accelerated phase or blast crisis, where RXFP1 expression patterns may differ [6, 23].

Additionally, the dataset does not provide information on patient demographics (e.g., age, sex) or clinical outcomes (e.g., response to therapy, survival rates), which could influence RXFP1 expression and its relevance as a biomarker [35]. Future studies should include a broader range of CML subtypes and incorporate clinical metadata to assess the applicability of RXFP1 as a biomarker across diverse patient populations [39].

### Probe-Level Redundancy in Biomarker Identification

A notable limitation in our biomarker identification process is the redundancy in probe-level expression data, particularly with PIEZO2. As discussed in Section 3.1, PIEZO2 is represented by three probe sets (8022295, 8022283, and 8022310) in the top 10 biomarker list, with importance scores of 0.1267, 0.1106, and 0.0778, respectively. This redundancy arises from several factors inherent to the dataset and preprocessing methodology, which are common issues in microarray studies [42, 48]:

- **Multiple Probe Sets**: PIEZO2 is represented by three distinct probe sets (8022295, 8022283, and 8022310), each targeting different regions of the gene (various segments on chromosome 18). This is a characteristic feature of the Affymetrix Human Gene 1.0 ST Array design, which employs multiple probe sets to cover different parts of a gene, ensuring comprehensive representation of its expression.
- **Transcript Variants**: PIEZO2 has multiple transcript variants, and each probe set targets a specific variant. Evidence from cDNA clones such as AK098782 and AK092226 supports the notion that these probe sets detect distinct splice variants of PIEZO2, contributing to its repeated representation in the dataset [10].
- **Probe Design and Annotation**: In the Affymetrix annotation, PIEZO2 is listed three times because each probe set targets a different segment or transcript of the gene. This is not a duplication error but rather an intentional aspect of the array’s design to capture the gene’s variability comprehensively.
- **Data Preprocessing**: During the data preprocessing stage, the probe sets were not summarized at the gene level. As a result, PIEZO2 appears as three separate entities associated with distinct probe set IDs, reflecting the lack of aggregation into a single gene-level representation. This redundancy highlights the overall importance of PIEZO2, potentially underestimating its biological significance in CML. Future studies should employ gene-level summarization to address this issue and improve the interpretability of biomarker rankings [21, 42].

## 8. Conclusion

This study identified RXFP1 as a robust biomarker for CML, characterized by consistent overexpression independent of BCR-ABL tyrosine kinase signaling. The lack of overlap between normal and CML expression values, even after imatinib treatment, underscores RXFP1’s diagnostic potential, a finding that aligns with other biomarker studies in CML [37, 43].

Furthermore, the independence of RXFP1 from BCR-ABL signaling suggests its potential involvement in BCR-ABL-independent mechanisms of TKI resistance, particularly in leukemia stem cells (LSCs) [8, 18]. However, the small sample size, potential experimental biases, and lack of functional validation necessitate further investigation. Future studies should focus on validating these findings in larger datasets, elucidating the regulatory mechanisms of RXFP1 overexpression, exploring its role in TKI resistance, and assessing its functional impact on LSC survival and CML pathogenesis [30, 45]. These findings open avenues for future investigations into the potential role of RXFP1 in influencing tyrosine kinase inhibitor (TKI) resistance and its association with achieving a deep molecular response (DMR) in CML patients, which we aim to explore in our subsequent experimental studies [19].

## 9. Data availability statement

The gene expression data analyzed in this study is publicly available at the Gene Expression Omnibus (GEO) under accession number GSE97562 (https://www.ncbi.nlm.nih.gov/geo/query/acc.cgi?acc=GSE97562). The data used for validation is publicly available at Gene Expression Omnibus (GEO) under accession number GSE279135 (https://www.ncbi.nlm.nih.gov/geo/query/acc.cgi?acc=GSE279135). The manual analysis summary of the GSE97562 dataset, detailing sample curation and organization into eight experimental groups, is provided in Supplementary File S1. Pathway enrichment analysis results, including p-values, q-values, source databases, and linked biomarkers, are provided in Supplementary Table S1. The detailed list of top 10 biomarkers identified by random forest modeling, including importance scores, AUC scores, p-values, and fold change values, is provided in Supplementary Table S2. All Python scripts, including the GUI tool and analysis pipelines, access restricted for now for security purpose will be openly accessible at Zenodo (DOI: 10.5281/zenodo.15575808).(Cross data validation included under research validation folder).

## 10. Funding

This research received no external funding.

## Supporting information

manual_analysis_GSE97562_summary

pathways_enrichment_tables

top 10 biomarkers table

supplimantry files description

## 11. Acknowledgments

The author acknowledges the support of [Your University/Institute Name] and the availability of the GSE97562 dataset on GEO.

## Notes

https://www.ncbi.nlm.nih.gov/geo/query/acc.cgi?acc=GSE97562

### Competing Interest Statement

The authors have declared no competing interest.

## References

1. Ahmed N, Yasin T, Elshoeibi AM, et al. Artificial intelligence-based management of adult chronic myeloid leukemia: Where are we and where are we going? J Pers Med. 2023;13(5):824.

2. Avilés-Vázquez S, Chávez-González A, Mayani H, et al. Global gene expression profiles of hematopoietic stem and progenitor cells from patients with chronic myeloid leukemia: the effect of in vitro culture with or without imatinib. Data Brief. 2017;14:576–582.

3. Baccarani M, Deininger MW, Rosti G, et al. European LeukemiaNet recommendations for the management of chronic myeloid leukemia: 2013. Blood. 2013;122(6):872–884.

4. Balboni S, Patergnani S, Giorgi C, et al. Relaxin/RXFP1 signaling in tumor biology: Possible opportunities for therapeutic intervention. Mol Cell Endocrinol. 2019;487:1–9.

5. Branford S, Rudzki Z, Walsh S, et al. High frequency of point mutations clustered within the adenosine triphosphate-binding region of BCR/ABL in patients with chronic myeloid leukemia or Ph-positive acute lymphoblastic leukemia who develop imatinib (STI571) resistance. Blood. 2002;99(9):3472–3475.

6. Breccia M, Celant S, Olimpieri PP, et al. Mortality rate in patients with chronic myeloid leukemia in chronic phase treated with frontline second-generation tyrosine kinase inhibitors: A retrospective analysis. Ann Hematol. 2021;100(2):481–485.

7. Calabrese G, Ferraresi A, Giandomenico S, et al. Resistance mechanisms to tyrosine kinase inhibitors in chronic myeloid leukemia: BCR-ABL dependent and independent pathways. Front Oncol. 2019;9:939.

8. Chomel JC, Turhan AG. Chronic myeloid leukemia stem cells: From basic science to clinical implications. Curr Hematol Malig Rep. 2015;10(4):389–397.

9. Corbin AS, Agarwal A, Loriaux M, et al. Human chronic myeloid leukemia stem cells are insensitive to imatinib despite inhibition of BCR-ABL activity. J Clin Invest. 2011;121(1):396–409.

10. Coste B, Mathur J, Schmidt M, et al. Piezo1 and Piezo2 are essential components of distinct mechanically activated ion channels. Science. 2010;330(6000):55-60.

11. Deininger MW, Goldman JM, Melo JV. The molecular biology of chronic myeloid leukemia. Blood. 2000;96(10):3343–3356.

12. Druker BJ, Talpaz M, Resta DJ, et al. Efficacy and safety of a specific inhibitor of the BCR-ABL tyrosine kinase in chronic myeloid leukemia. N Engl J Med. 2001;344(14):1031–1037.

13. Eckardt JN, Bornhäuser M, Wendt K, et al. Application of machine learning in the management of chronic myeloid leukemia: Current practice and future prospects. Blood Adv. 2021;5(23):6077–6085.

14. Elsabagh AA, Mohamed RA, Loza NF, et al. Application of artificial intelligence in chronic myeloid leukemia (CML) disease prediction and management: A scoping review. BMC Cancer. 2024;24(1):1002.

15. Feng S, Agoulnik IU, Bogatcheva NV, et al. Relaxin promotes prostate cancer progression. Clin Cancer Res. 2010;16(5):1491–1500.

16. Graham SM, Jørgensen HG, Allan E, et al. Primitive, quiescent, Philadelphia-positive cells from patients with chronic myeloid leukemia are insensitive to STI571 in vitro. Blood. 2002;99(1):319–325.

17. Halls ML, Bathgate RA, Sutton SW, et al. International Union of Basic and Clinical Pharmacology. XCV. Recent advances in the understanding of the pharmacology and biological roles of relaxin family peptide receptors 1–4, the receptors for relaxin family peptides. Pharmacol Rev. 2015;67(1):1–25.

18. Hamilton A, Helgason GV, Schemionek M, et al. Chronic myeloid leukemia stem cells are not dependent on BCR-ABL kinase activity for their survival. Blood. 2012;119(6):1501–1510.

19. Hehlmann R, Müller MC, Lauseker M, et al. Deep molecular response is reached by the majority of patients treated with imatinib, predicts survival, and is achieved more quickly by optimized high-dose imatinib: Results from the randomized CML-Study IV. J Clin Oncol. 2014;32(5):415–423.

20. Hochhaus A, Larson RA, Guilhot F, et al. Long-term outcomes of imatinib treatment for chronic myeloid leukemia. N Engl J Med. 2017;376(10):917–927.

21. Huang DW, Sherman BT, Lempicki RA. Systematic and integrative analysis of large gene lists using DAVID bioinformatics resources. Nat Protoc. 2009;4(1):44–57.

22. Hunter JD. Matplotlib: A 2D graphics environment. Comput Sci Eng. 2007;9(3):90–95.

23. Jabbour E, Kantarjian H. Chronic myeloid leukemia: 2020 update on diagnosis, therapy, and monitoring. Am J Hematol. 2020;95(6):691-709.

24. Jassal B, Matthews L, Viteri G, et al. The Reactome pathway knowledgebase. Nucleic Acids Res. 2020;48(D1):D498–D503.

25. Kamimura K, Yokoo T, Fujimoto Y, et al. RXFP1 as a novel biomarker for liver fibrosis in patients with non-alcoholic fatty liver disease. Hepatol Res. 2020;50(6):698–707.

26. Kanehisa M, Goto S. KEGG: Kyoto encyclopedia of genes and genomes. Nucleic Acids Res. 2000;28(1):27–30.

27. Liberzon A, Subramanian A, Pinchback R, et al. Molecular signatures database (MSigDB) 3.0. Bioinformatics. 2011;27(12):1739–1740.

28. McKinney W. Data structures for statistical computing in Python. Proc 9th Python Sci Conf. 2010;445:51–56.

29. McWeeney SK, Pemberton LC, Loriaux MM, et al. A gene expression signature of CD34+ cells to predict major cytogenetic response in chronic-phase chronic myeloid leukemia patients treated with imatinib. Blood. 2010;115(2):315–325.

30. Neviani P, Harb JG, Oaks JJ, et al. PP2A-activating drugs selectively eradicate TKI-resistant chronic myeloid leukemic stem cells. J Clin Invest. 2013;123(10):4144–4157

31. . Ni J, Wang X, Li Y, et al. A novel flow cytometry method using support vector machine for distinguishing malignant from normal neutrophils in chronic myeloid leukemia. Cytometry B Clin Cytom. 2021;100(3):312–320.

32. O’Brien SG, Guilhot F, Larson RA, et al. Imatinib compared with interferon and low-dose cytarabine for newly diagnosed chronic-phase chronic myeloid leukemia. N Engl J Med. 2003;348(11):994–1004.

33. Oehler VG, Yeung KY, Choi YE, et al. The derivation of diagnostic markers of chronic myeloid leukemia progression from microarray data. Blood. 2009;114(15):3292–3298.

34. Pedregosa F, Varoquaux G, Gramfort A, et al. Scikit-learn: Machine learning in Python. J Mach Learn Res. 2011;12:2825–2830.

35. Pfirrmann M, Baccarani M, Saussele S, et al. Prognosis of long-term survival considering disease-specific death in patients with chronic myeloid leukemia. Leukemia. 2016;30(1):48–56.

36. Radich JP, Dai H, Mao M, et al. Gene expression changes associated with progression and response in chronic myeloid leukemia. Proc Natl Acad Sci U S A. 2006;103(8):2794–2799.

37. Raspadori D, Pacelli P, Sicuranza A, et al. Flow cytometry assessment of CD26(+) leukemic stem cells in peripheral blood: A simple and rapid new diagnostic tool for chronic myeloid leukemia. Cytometry B Clin Cytom. 2019;96(4):294–299.

38. Samuel CS, Royce SG, Hewitson TD, et al. Anti-fibrotic actions of relaxin: From bench to bedside. Br J Pharmacol. 2017;174(10):969–985.

39. Sasaki K, Jabbour E, Ravandi F, et al. The impact of treatment recommendation by leukemia artificial intelligence program (LEAP) on survival in patients with chronic myeloid leukemia in chronic phase (CML-CP). Blood. 2019;134(Suppl_1):130148.

40. Singh S, Bennett RG. Relaxin family peptide receptors (RXFP1-4): Structure, function, and signaling. Front Endocrinol (Lausanne). 2021;12:691802.

41. Stagno F, Russo S, Murdaca G, et al. Utilization of machine learning in the prediction, diagnosis, prognosis, and management of chronic myeloid leukemia. Int J Mol Sci. 2025;26(6):2535.

42. Subramanian A, Tamayo P, Mootha VK, et al. Gene set enrichment analysis: A knowledge-based approach for interpreting genome-wide expression profiles. Proc Natl Acad Sci U S A. 2005;102(43):15545–15550.

43. Sweet K, Zhang L, Pinilla-Ibarz J. Biomarkers for determining the prognosis in chronic myelogenous leukemia. J Hematol Oncol. 2013;6:54.

44. Thompson VC, Morris TG, Cochrane DR, et al. Relaxin and estrogen synergize to promote breast cancer progression via RXFP1 signaling. Mol Cancer Res. 2016;14(8):703–714.

45. Tyner JW, Tognon CE, Bottomly D, et al. Functional genomic landscape of acute myeloid leukemia. Nature. 2018;562(7728):526-531.

46. Van Rossum G, Drake FL. Python 3 reference manual. CreateSpace; 2009.

47. Waskom M. Seaborn: Statistical data visualization. J Open Source Softw. 2021;6(60):3021.

48. Yeoh EJ, Ross ME, Shurtleff SA, et al. Classification, subtype discovery, and prediction of outcome in pediatric acute lymphoblastic leukemia by gene expression profiling. Cancer Cell. 2002;1(2):133–143.

49. Zhang Y, Li Z, Zhang J, et al. Automatic diagnosis of chronic myeloid leukemia using conditional generative adversarial networks. Am J Pathol. 2022;192(5):732–742.

50. Zhong Z, Zhang H, Li X, et al. Identification of diagnostic biomarkers for chronic myeloid leukemia using machine learning approaches. Front Oncol. 2022;12:852345.

51. Biernaux, C., et al. (1995). Detection of major bcr-abl gene expression at a very low level in blood cells of some healthy individuals. Blood, 86(2), 630–636.

52. Bayraktar, S., & Goodman, M. (2010). Asymptomatic BCR-ABL-positive individuals: a review of implications for CML pathogenesis. Leukemia Research, 34(11), 1432–1436.

