## Supplementary material for "AI-Driven Identification and Validation of RXFP1 as a CML Biomarker Using Gene Expression and Integrated GUI Tools": manual_analysis_GSE97562_summary

### Manual Analysis Summary of GSE97562_series_matrix Data

A total of 40 gene expression samples were curated and organized into 8 distinct experimental groups (5 samples per group), based on disease status (CML vs. normal), cell lineage (stem vs. progenitor), culture time (T0 vs. 48 hours), and drug exposure (± imatinib). The classification is detailed as follows:

1. Indices 0-4:

Gene expression data of chronic myeloid leukemia (CML) stem cells at time point T0, isolated using CD34 and CD38 markers via flow cytometry.

(Untreated CML stem cells at baseline)

2. Indices 5-9:

Gene expression data of normal bone marrow stem cells at time point T0, isolated using CD34 and CD38 markers via flow cytometry.

(Untreated normal stem cells at baseline)

3. Indices 10-14:

Gene expression data of CML progenitor cells at time point T0, isolated using CD34 and CD38 markers via flow cytometry.

(Untreated CML progenitor cells at baseline)

4. Indices 15-19:

Gene expression data of normal bone marrow progenitor cells at time point T0, isolated using CD34 and CD38 markers via flow cytometry.

(Untreated normal progenitor cells at baseline)

5. Indices 20-24:

Gene expression data of CML progenitor cells cultured for 48 hours without imatinib, isolated using CD34 and CD38 markers via flow cytometry.

(CML progenitor cells post 48-hour culture without treatment)

6. Indices 25-29:

Gene expression data of normal progenitor cells cultured for 48 hours without imatinib, isolated using CD34 and CD38 markers via flow cytometry.

(Normal progenitor cells post 48-hour culture without treatment)

7. Indices 30-34:

Gene expression data of CML progenitor cells cultured for 48 hours with imatinib, isolated using CD34 and CD38 markers via flow cytometry.

(CML progenitor cells post 48-hour culture with imatinib treatment)

8. Indices 35-39:

Gene expression data of normal progenitor cells cultured for 48 hours with imatinib, isolated using CD34 and CD38 markers via flow cytometry.

(Normal progenitor cells post 48-hour culture with imatinib treatment)

This experimental design enables comprehensive comparative transcriptomic analysis across multiple biological dimensions-disease progression, cell differentiation, time-course response, and pharmacological intervention. Such stratification ensures reliable insight into gene regulation dynamics in the context of chronic myeloid leukemia and normal hematopoiesis.
