## Supplementary material for "AI-Driven Identification and Validation of RXFP1 as a CML Biomarker Using Gene Expression and Integrated GUI Tools": pathways_enrichment_tables

| Pathway | p-value | q-value | Source | Linked Biomarker |
|---|---|---|---|---|
| Relaxin Receptors | 0.002797 | 0.040882 | BioPlanet | RXFP1 |
| Tryptophan Catabolism | 0.003844 | 0.040882 | BioPlanet | KYNU |
| Caspase-mediated Cleavage of Cytoskeletal Proteins | 0.004193 | 0.040882 | BioPlanet | GAS2 |
| N-cadherin Signaling Events | 0.011841 | 0.048540 | BioPlanet | CDH2 |
| Adherens Junctions Interactions | 0.010107 | 0.048540 | BioPlanet | CDH2 |
| Voltage Gated Potassium Channels | 0.014955 | 0.048540 | BioPlanet | KCNA3 |
| Apoptotic Execution Phase | 0.018061 | 0.048540 | BioPlanet | GAS2 |
| Tryptophan Metabolism | 0.020127 | 0.048540 | BioPlanet | KYNU |
| Caspase Cascade in Apoptosis | 0.020471 | 0.048540 | BioPlanet | GAS2 |
| Post-translational Protein Phosphorylation | 0.036859 | 0.095047 | Reactome | (CML-related) |
