## Supplementary material for "AI-Driven Identification and Validation of RXFP1 as a CML Biomarker Using Gene Expression and Integrated GUI Tools": top 10 biomarkers table

**Table 1: Top 10 Biomarkers for Chronic Myeloid Leukemia (CML) Detection**

This table presents the top 10 biomarkers identified for CML detection using a Random Forest classifier, based on gene expression data. The biomarkers are ranked by their importance scores, reflecting their contribution to distinguishing CML from normal samples. The table includes differential expression metrics (log2FC, p-value, -log10(p_value)) to highlight the statistical significance of each biomarker's expression difference between CML and normal samples.

| **Rank** | **Gene ID** | **Gene Symbol** | **Gene Name** | **log2FC** | **p-value** | **-log10(p_value)** | **Importance Score** |
| --- | --- | --- | --- | --- | --- | --- | --- |
| 1 | 8098060 | RXFP1 | Homo sapiens relaxin/insulin like family peptide receptor 1 (RXFP1) | 0.4628 | 2.0450e-12 | 11.69 | 0.1393 |
| 2 | 8022295 | PIEZO2 | Homo sapiens piezo type mechanosensitive ion channel component 2 (PIEZO2) | 0.4307 | 4.1696e-14 | 13.38 | 0.1267 |
| 3 | 8022283 | PIEZO2 | Homo sapiens piezo type mechanosensitive ion channel component 2 (PIEZO2) | 0.5198 | 2.6155e-12 | 11.58 | 0.1106 |
| 4 | 8022310 | PIEZO2 | Homo sapiens piezo type mechanosensitive ion channel component 2 (PIEZO2) | 0.4373 | 3.7309e-12 | 11.43 | 0.0778 |
| 5 | 8022674 | CDH2 | Homo sapiens cadherin 2 (CDH2), transcript variant 1, mRNA | -0.3155 | 8.3816e-08 | 7.08 | 0.0731 |
| 6 | 7938989 | GAS2 | Homo sapiens growth arrest specific 2 (GAS2), transcript variant 3, mRNA | 0.4132 | 4.4704e-04 | 3.35 | 0.0513 |
| 7 | 7918457 | KCNA3 | Homo sapiens potassium voltage-gated channel subfamily A member 3 (KCNA3) | -0.3896 | 2.4859e-07 | 6.60 | 0.0489 |
| 8 | 8095819 | FAM47E-STBD1 | Homo sapiens family with sequence similarity 47 member E (FAM47E) | 0.3947 | 8.0747e-11 | 10.09 | 0.0409 |
| 9 | 8045539 | KYNU | Homo sapiens kynureninase (KYNU), transcript variant 2, mRNA | 0.3017 | 1.5215e-07 | 6.82 | 0.0389 |
| 10 | 7902102 | Unknown | Not Available | 0.3810 | 1.3551e-06 | 5.87 | 0.0353 |

**Notes:**

- **Rank:** Biomarkers are ranked based on their importance scores from the Random Forest model.
- **Gene ID:** Unique identifier for each gene in the dataset.
- **Gene Symbol:** Official gene symbol as per the Human Genome Organization (HUGO) nomenclature. Gene ID 7902102 lacks a corresponding symbol in the provided data and is marked as "Unknown."
- **Gene Name:** Official name or description of the gene, extracted from the nucleotide title. For Gene ID 7902102, the gene name is marked as "Not Available" due to missing data.
- **log2FC:** Log2 fold change of gene expression between CML and normal samples (positive values indicate higher expression in CML; negative values indicate lower expression). Rounded to 4 decimal places.
- **p-value:** Statistical significance of the differential expression, calculated using a suitable statistical test (e.g., t-test or Mann-Whitney U test). Presented in scientific notation.
- **-log10(p_value):** Negative log10 transformation of the p-value, indicating the strength of statistical significance (higher values denote greater significance). Rounded to 2 decimal places.
- **Importance Score:** Contribution of each gene to the Random Forest model's predictive performance for CML detection, averaged across 5-fold cross-validation. Rounded to 4 decimal places.
- **Data Source:** Gene expression data was derived from the processed dataset (expression_with_labels.csv), and importance scores were computed using a Random Forest classifier trained on 35 selected genes. Gene symbols and names were mapped from the provided metadata.
